# The efficiency of p27 peptide cleavage during in vitro respiratory syncytial virus (RSV) infection is cell line and RSV subtype dependent

**DOI:** 10.1101/2022.10.17.512554

**Authors:** Wanderson Rezende, Xunyan Ye, Laura S. Angelo, Alexandre Carisey, Vasanthi Avadhanula, Pedro A. Piedra

## Abstract

Respiratory Syncytial Virus (RSV) Fusion protein (F) is highly conserved between RSV/A and RSV/B subtypes. To become fully active, F precursor undergoes enzymatic cleavage to yield F1 and F2 subunits and releases a 27 amino acid peptide (p27). Virus-cell fusion occurs when RSV F undergoes a conformational change from pre-F to post-F. Previous immunological and cell-surface expression data show that p27 is detected on RSV F, but questions remain on how p27 effects the conformation of mature RSV F. Monoclonal antibodies against p27, Site Ø (pre-Fusion specific), and Site II were used to monitor RSV F conformation by ELISA and Imaging Flow Cytometry. Pre-F to post-F conformational change was induced by temperature-stress test. We found that p27 cleavage efficiency was lower on sucrose purified (sp) RSV/A than on spRSV/B. In addition, *in vitro* cleavage of RSV F was cell-line dependent, higher levels of p27 expression were observed on surface of RSV infected HEp-2 cells than A549 cells. Higher levels of p27 were also found on RSV/A infected cells compared to RSV/B. We observed that RSV/A F with higher levels of p27 could better sustain the pre-F conformation during the temperature-stress challenge in both spRSV and as well RSV-infected cell lines. Our findings suggest that despite F sequence similarity, the p27 of RSV subtypes is cleaved with different efficiencies, which were also dependent on the cell lines used for infection. We therefore speculate that partially cleaved p27 may confer higher stability to the pre-F and provides a fitness advantage.

**IMPORTANCE:** The RSV fusion protein (F) plays an important role in entry and viral fusion to the host cell. The F protein undergoes proteolytic cleavages by furin protease resulting in the release of a 27 amino acid peptide (p27) to become fully functional. During this process, the F protein also undergoes a conformational change from metastable pre-F to highly stable post-F. For decades, the consensus in the RSV field was that p27 was not present in the fully mature RSV F protein. Therefore, the role of the p27 peptide in viral entry and the function of the partially cleaved F protein containing p27 has been overlooked. However, recent developments have shown that p27 elicits an immune response during natural infection and that p27 can be found *in vitro* and in animal models infected with the prototypical RSV/A strain. In this study, we were able to detect p27 on purified RSV virions and on the surface of virus-infected cells of two widely used cell lines, HEp-2 and A549 cells for both prototypical and contemporary circulating RSV strains of both subtypes. Also, higher levels of partially cleaved F protein containing p27 could better sustain the pre-F conformation during the temperature-stress challenge. Our findings highlight that the cleavage efficiency of p27 is different between RSV subtypes and among cell lines and that the presence of p27 in partially cleaved F protein likely contributes to the stability of the pre-F conformation.

## INTRODUCTION

Respiratory Syncytial Virus (RSV) is one of the major causes of acute lower respiratory infections (ALRI), imposing a significant impact on global public health (1–4). RSV primarily impacts children under the age of five years, older adults, and immunocompromised individuals (5–8), but all humans experience RSV reinfections throughout life (9). Prior to the SARS-CoV-2 pandemic, RSV outbreaks occurred predictably during the fall and winter months in temperate climates (10). Generally, during an outbreak, one of the two subtypes of RSV (RSV/A and RSV/B) predominates (11, 12). RSV subtypes are further subdivided into genotypes that can co-circulate at a given time (11, 13). Since 2011 and 2014, RSV/A Ontario (ON) and RSV/B Buenos Aires (BA) have been the two predominant circulating RSV genotypes around the globe (12).

RSV is an enveloped, negative-sense RNA virus belonging to the *Pneumoviridae* family. Of the 11 proteins encoded by its 15.2 kilobase genome, three are located on the viral envelope: the attachment (G), small hydrophobic (SH), and the fusion proteins (F) (14, 15). Epitope differences in the G protein – and to a lesser degree in the F protein – divide RSV into subtypes RSV/A and RSV/B (13, 14). Interestingly, however, the major target of vaccines and monoclonal antibodies is the fusion (F) protein of RSV. The F protein is highly conserved between RSV/A and RSV/B (16, 17), making it an excellent target for therapeutic development. It is a Class I transmembrane glycoprotein synthesized as a precursor (F0), becoming active, or fusogenic, upon enzymatic cleavage of an internal peptide of 27 amino acids (p27) between RARR109 furin cleavage site 2 (FCS-2) and KKRKRR136 (FCS-1). The cleavage of F0 occurs intracellularly by furin-like enzymes, which yields two subunits of distinct sizes: F1 (50 kDa) and F2 (20 kDa) with the release of p27 peptide (18, 19).

The F protein plays a vital role in the early phase of infection, allowing the fusion of the virus membrane to the host cell membrane (15, 20). The RSV F protein can retain p27 by undergoing partial cleavage, as cleavage on FCS-2 is dispensable for viral fitness (21, 22) and can also exist as uncleaved F0 on the surface of RSV viruses and infected cells (22). This is relevant as there are two concurrent observations pertaining to the mechanism of RSV fusion and entry into the host cell. The first proposes that the F proteins on infectious RSV are fully cleaved (no p27), and RSV fuses directly with the cell membrane. The second mechanism proposes that the F protein on RSV harbors partially cleaved p27 and it is internalized through actin-dependent micropinocytosis; a second enzymatic cleavage excises the remaining p27 and the RSV gains access to the cytoplasm by fusing with the endosomal membrane (22). Thus, the relationship between p27 cleavage and F protein is central to all proposed mechanisms of RSV entry in host cells.

The biological role of p27 has been gaining interest in the scientific community as we and others observed that p27 elicits a strong immune response in RSV-infected children and adults by both RSV/A and RSV/B (23–25), thus supporting the likelihood of partially cleaved F containing p27 during viral infection. A recent study showed that p27 can be detected on the surface of RSV-infected cells and the lungs of infected mice (26), suggesting that p27 is broadly present in RSV-infected cells *in vitro* and *in vivo*. Even though groundbreaking studies using X-ray crystallography and Cryo-EM spectroscopy have improved our understanding of the conformation-dependent function of the RSV F protein (27–30), to this day, no structure containing p27 has been generated, mostly due to the disordered nature of the region and to the low expression yields from recombinant constructs containing the peptide (31–33). To date, some limitations of the structural studies were the use of prototypical strains of RSV/A (nowadays far removed from circulating contemporary strains) and the need for structural modifications that stabilize the metastable pre-F conformation. Such strategies were crucial to generating high-resolution structures and led to identifying conformation-specific monoclonal antibodies targeting RSV F protein epitopes.

Structural studies coupled with immunoassays using epitope-specific monoclonal antibodies showed that RSV F protein has a dynamic quaternary structure, existing in equilibrium between monomer and trimer (34). As a trimer, the F protein contains at least three antigenic sites (sites Ø, III & V) unique to the metastable pre-F conformation. Once triggered, the pre-F undergoes an irreversible rapid conformational transition to a highly stable post-F form (35). The post-F contains two antigenic sites that are shared with the pre-F form (sites II and IV) and at least one antigenic Site unique to post-F (Site I). Due to the metastability of the pre-F, the conformational change may also happen spontaneously, hindering the F protein of its fusogenic function (36). The mechanism and factors that drive this structural rearrangement are not well understood, but the conformation rearrangement must happen near host cell membranes (via G and F proteins) for productive fusion (37).

To explain both the immunological observation with p27 and the instability of pre-F conformation, we hypothesize that RSV F can exist as a partially cleaved protein, retaining p27 (pre-F_p27_) in infectious virions. We also hypothesize that the presence of p27 helps stabilize the pre-F conformation. We utilized monoclonal antibodies (mAb) specific to p27 (38), Site Ø (unique to pre-F), and Site II (shared by pre-F and post-F) (33, 39) to study the presence of p27 on infectious virions as well as on cell surface of virus-infected cells and used temperature-stress studies to evaluate pre-F stability.

Our findings demonstrate that p27 is present on infectious virions as well as on the surface of RSV-infected cells and is mostly associated with pre-F conformation. A comparison of both prototypic and contemporary RSV/A and B isolates revealed that RSV/A isolates contain a higher proportion of p27 on their F protein. We also observed a host effect on the cleavage efficiency of p27, whereby RSV/A or RSV/B-infected HEp-2 cells retained a greater number of virus-infected cells containing pre-F_p27_ as compared to RSV-infected A549 cells of either subtype. Lastly, using a temperature-stress test to trigger pre-F to post-F conformation, the presence of p27 appeared to be associated with improvement of pre-F stability with elevated temperatures in these physiological contexts. Overall, our findings show that p27 is commonly detected in the F proteins of RSV/A and RSV/B strains, and viral and host factors play a central role in the generation of partially cleaved F that retains the p27 with greater stability. These observations support the role of pre-F_p27_ in RSV infection.

## MATERIALS AND METHODS

### Cells, viruses, and antibodies

Human epidermoid carcinoma larynx cell line (HEp-2, ATCC) and adenocarcinomic human alveolar basal epithelial cells (A549, ATCC) were cultured at 36 °C (5% CO_2_) in complete MEM (Modified Eagle’s medium (MEM, Corning 10-010-CM) supplemented with 10% fetal bovine serum (Hyclone SH30070.03), 2 mM L-glutamine (Gibco 25030081), and antibiotic-antimycotic (100 U/ml penicillin/streptomycin/ 0.25µg/mL Amphotericin B (Gibco 15240062)).

Working pools of RSV/A/USA/BCM-Tracy/1989 (genotype GA1), RSV/A/WashingtonDC.USA/Bernett/1961 (genotype GA1), RSV/B/WashingtonDC.USA/18537/1962 (genotype GB1), RSV/A/USA/BCM813013/2013 (genotype ON), and RSV/B/USA/BCM80171/2010 (genotype BA) will be referred to as RSV/A/Tracy, RSV/A/Bernett, RSV/B/18537, RSV/A/ON and RSV/B/BA, respectively. RSV/A/Bernett and RSV/B/BA were purified by ultracentrifugation on a sucrose cushion (spRSV) as previously described (40, 41). RSV/A/Bernett was the virus strain used in the 1960s to develop the formalin-inactivated RSV vaccine that was associated with vaccine-enhanced disease in infants (42). RSV/A/Tracy, RSV/A/Bernett and RSV/B/18537 are prototypic strains while RSV/A/ON and RSV/B/BA are contemporary strains.

RSV F specific monoclonal antibodies (mAbs) against antigenic Sites Ø (D25, Cambridge Biologics, LLC, Brookline, MA, USA), Site II (Palivizumab^®^, MedImmune, LLC, Geithersburg, MD), and p27 (RSV7.10, kindly provided by Dr. Gale Smith (Novavax, MD)) were used to assign the conformation of the F protein and to determine the presence of p27. Pre-F conformation was assigned by the detection of Sites Ø and II. Post-F conformation was assigned by the absence of Site Ø and the presence of Site II.

Horseradish peroxidase (HRP)-conjugated goat anti-mouse IgG (BioRad #172-1011) and HRP-conjugated goat anti-human IgG (BioRad #172-1050) were used as secondary antibodies for ELISA and western blot assays.

### Western Blot

The total protein concentration on sucrose-purified RSV/A/Bernett and RSV/B/BA was determined by micro-BCA assay following the manufacturer protocol (ThermoScientific, 23235). Approximately 10 μg of each spRSV was mixed with SDS-Laemmli buffer under reducing conditions (50 μm of BME) and incubated at 95°C for 5 minutes. The samples were run on a 4– 20% Mini-PROTEAN TGX Precast Protein Gels (Bio-Rad, 456-1095) at 200V for 25 minutes, then transferred to a nitrocellulose membrane by semi-dry technique (Trans-Blot Turbo Transfer System, Bio-Rad, 1704150). The membrane was blocked with 1% Casein in TBS (Bio-Rad, 161-0782) for one hour at room temperature and incubated with D25, Palivizumab, or RSV7.10 mAb (anti-antigenic sites Ø, II, and p27, respectively) diluted in blocking buffer (1:1,000, 1:4000, and 1:1000, respectively) for one hour at room temperature and washed three times with PBST (0.1%-Tween 20). Membranes were then incubated for one hour at room temperature with HRP-conjugated goat anti-human IgG (membranes probed with D25 or Palivizumab) or HRP-conjugated goat anti-mouse IgG (membranes probed with RSV7.10) at 1:2000 dilution in PBST. After four washes with PBST, the bands were developed with Clarity Western ECL Substrate (Bio-Rad, 170-5060) and membranes imaged with a ChemiDoc MP System (Bio-Rad, 12003154).

### Enzyme-Linked Immunosorbent Assays (ELISA)

The relative proportions of Site Ø, Site II, and p27 in spRSVs were determined by a modified ELISA protocol from Ye et al. (43). In short, 96-well microtiter plates (Immulon 2HB plates, Thermo Scientific) were coated with sequentially diluted spRSV/A/Bernett or spRSV/B/BA in PBS (starting at 10 µg/mL or 15 µg/mL of total protein, respectively) for 18-20 hours at 4 °C. PBS at pH 7.2 was used as coating buffer instead of carbonate/bicarbonate buffer at pH 9.6 as it preserved the binding of D25 to the pre-F conformation. Plates were then washed three times with 1X Kirkegaard and Perry Laboratories (KPL) wash buffer (SeraCare Life Sciences, Gaithersburg, MD), and blocked with 5% milk (Carnation Instant Nonfat Dry Milk) in KPL washing buffer for one hour at 36 °C. Monoclonal antibodies D25, Palivizumab, and RSV7.10 (anti-antigenic sites Ø, II, and p27, respectively) were diluted at 1.1 µg/mL in blocking buffer, added to the plate and incubated for one hour at 36 °C. After three washes with KPL washing buffer, HRP-conjugated secondary antibodies goat anti-human IgG or goat anti-mouse IgG were diluted at 1:2000 in KPL washing buffer and added to the plates for one hour at 36 °C. After six washes with KPL buffer, the signal was developed with 3,3’,5,5’-Tetramethylbenzidine (TMB, Cat. # 50-76-03, Kirkegaard and Perry Labs) for 18 minutes in the dark at room temperature. Developing reaction was stopped with 0.16 M sulfuric acid and plates were promptly read at 450 nm wavelength on a BioTek Synergy H1 microplate reader.

### Temperature stress test to evaluate the stability of pre-F conformation on infectious sucrose-purified RSV

spRSV/A/Bernett or spRSV/B/BA at total protein concentration of 1.0 mg/mL and 0.72 mg/mL, respectively, were diluted five-fold in distilled water, and aliquots heated at various temperatures (25°C, 30°C, 40°C, 50°C, 60°C, 70°C, 80°C, and 90 °C) for 10 minutes in a digital heat block. After each temperature treatment, samples were cooled to 25 °C and diluted 16-fold in PBS. The wells of 96-well microtiter plates (Immulon 2HB plates, Thermo Scientific) were coated with 100 µL of each diluted aliquot in triplicate and incubated for 18-20 hours at 4 °C. ELISA protocol was then executed as described above.

### Imaging Flow Cytometry

Fluorescently tagged antibodies were generated by covalently labeling monoclonal antibodies D25 (Site Ø), Palivizumab (Site II) and RSV7.10 (p27) with AlexaFluor 568, AlexaFluor 488, and AlexaFluor 647, respectively, according to the manufacturer’s protocol (Invitrogen, Cat# A20184, A20181, and A20186).

HEp-2 and A549 cells were cultured to confluence as described above and infected at an MOI of0.07. At each determined time point – Days 1, 2, 3, 4 and 5 post-inoculation – the cells were washed with PBS and detached with Versene solution for 20 minutes at 36 °C (0.48 mM EDTA in PBS; ThermoFisher, Cat# A4239101). The cells (10^5^ – 10^6^ cells per staining condition) were resuspended in FACS Buffer (2% BSA in PBS, sterile), blocked with Fc Block for 20 min at room temperature (Human TruStain FcX, Biolegend Cat# 422302), and simultaneously stained for surface antigenic Site Ø, Site II, and p27 with primary-conjugated antibodies (D25-AF568, Palivizumab-AF488, and RSV7.10-AF647 at 8 µg/µL, respectively) for 30 min on ice. After washing with FACS Buffer, cells were stained with Hoechst 33342 at 5 ng/µL (ThermoFisher, Cat# H3570) for 10 min on ice. Cells were washed with PBS and fixed in 4% PFA, followed by acquisition by an Amnis Mark II ImageStream (405 nm, 488 nm, 561 nm, and 642 nm excitation lasers, 40× magnification) using the INSPIRE software (Luminex). Using the brightfield channel, a cell area vs cell aspect ratio plot was used to gate single cells from beads and cellular debris. Subsequently, out of focus objects were gated out from the dataset using the gradient RMS measurement (<45%) on the same brightfield channel. For all RSVs, the same number of events were collected at each day post-inoculation (dpi) based on the maximum number of cells at each dpi remaining attached to the culture dish (Supplemental Figure 1) (10,000 cells at days 1 to 3; 1,000 cells at day 4; 500 cells at day 5). Single-stained cells were used for spectral compensation, while uninfected cells and fluorescence-minus-one (FMO) controls were used for gating strategy. Data analysis was performed using IDEAS software.

### Temperature stress test to evaluate the stability of pre-F conformation on RSV-infected HEp-2 or A549 cells

HEp-2 and A549 cells were cultured to confluence as described above and infected with RSV at an MOI of 0.07. At 72h post-inoculation, the cells were washed with PBS and detached with Versene solution for 20 minutes at 36 °C (0.48 mM EDTA in PBS; ThermoFisher, Cat# A4239101). Next, the cells were resuspended in PBS at 10^6^ cells/400 µL aliquots and heated at various temperatures (36°C, 45°C, 50°C, 55°C, 60°C, and 65°C) for 10 minutes in a digital heat block. Cells were kept at 4 °C until ready to stain with the same protocol as described above. After fixation, cells were analyzed on Amnis Mark II ImageStream (405 nm, 488 nm, 561 nm, and 642 nm excitation lasers, 40× magnification; EMD-Millipore) and data analysis was performed using IDEAS software (Amnis).

### Imaging flow cytometry data analysis

Fluorescence intensity and image analysis were performed with the Amnis IDEAS software (Luminex), with all gate values determined empirically using unstained cells and FMO controls from each condition. Gating strategy is shown in Supplemental Figure 1. In short, in-focus cells were gated by an FSC gradient root-mean-square (RMS) histogram, followed by the selection of events containing one object per image (cell count mask). Nucleated and multinucleated cells (syncytia) were gated using a custom nuclei mask. A mask was created to isolate the fluorescence intensity signal from the cell membrane using the brightfield reference image to minimize the contribution of intracellular and nuclear background fluorescence. The percentage of RSV-infected cells (Site II-positive events) expressing F proteins on the pre-F conformation (Site Ø-positive events) were gated using a bivariate plot of pixel intensity for Site Ø (568 nm) vs. Site II (488 nm). The percentage of RSV-infected cells with F proteins containing p27 was gated by a bivariate plot of pixel intensity for p27 (647 nm) vs. Site II (488 nm).

## RESULTS

### Peptide 27 detected by Western blot

The prevailing dogma puts forth that infectious RSV viruses only contain fully cleaved F protein primarily in the pre-F conformation, without p27. Our RSV immunogenicity data from natural infection are not consistent with this point of view (23, 25). To determine if infectious RSV virions retained p27 on its F protein, we probed spRSV/A/Bernett (Fig. 1A) and spRSV/B/BA (Fig. 1B) for Site II (Palivizumab, Synagis), Site Ø (D25, Cambridge Bio), or p27 (RSV7.10) under reducing and denatured conditions by Western blot. Binding with Palivizumab of spRSV/A/Bernett revealed a 50 kDa band corresponding to the F1 subunit, and a 60 kDa band corresponding to the F1 subunit retaining a partially cleaved p27 peptide (F1+p27) (Fig. 1A, Site II). The bands on the nitrocellulose membrane probed with the p27 mAb confirmed the presence of the F1+p27 species (∼60 kDa), and a 70 kDa band likely corresponding to an uncleaved F protein (F0), which was not detected by either Palivizumab or D25 mAbs. Interestingly, D25 also detected a 60 kDa band likely corresponding to the F1+p27 species. The Western blot assay of spRSV/B/BA (Fig. 1B) showed a different distribution of F protein species. Binding with Palivizumab and p27 mAbs revealed only a single 60 kDa band corresponding to the partially cleaved F1+ p27 species, whereas the band corresponding to the completely cleaved F1 subunit (50 kDa) was not detected. Similar to the finding for RSV/A/Bernett, D25 mAb detected a 60 kDa band in the spRSV/B/BA. These findings show that p27 mAb detected a 60 kDa band in both RSV/A and RSV/B strains consistent with a partially cleaved F protein containing p27. In addition, the presence of uncleaved F protein (F0, 70 kDa) and fully cleaved F1 (50 kDa band) on spRSV/A/Bernett but not on spRSV/B/BA suggests that furin cleavage efficiency between RSV subtypes can differ (22).

**Figure 1:**
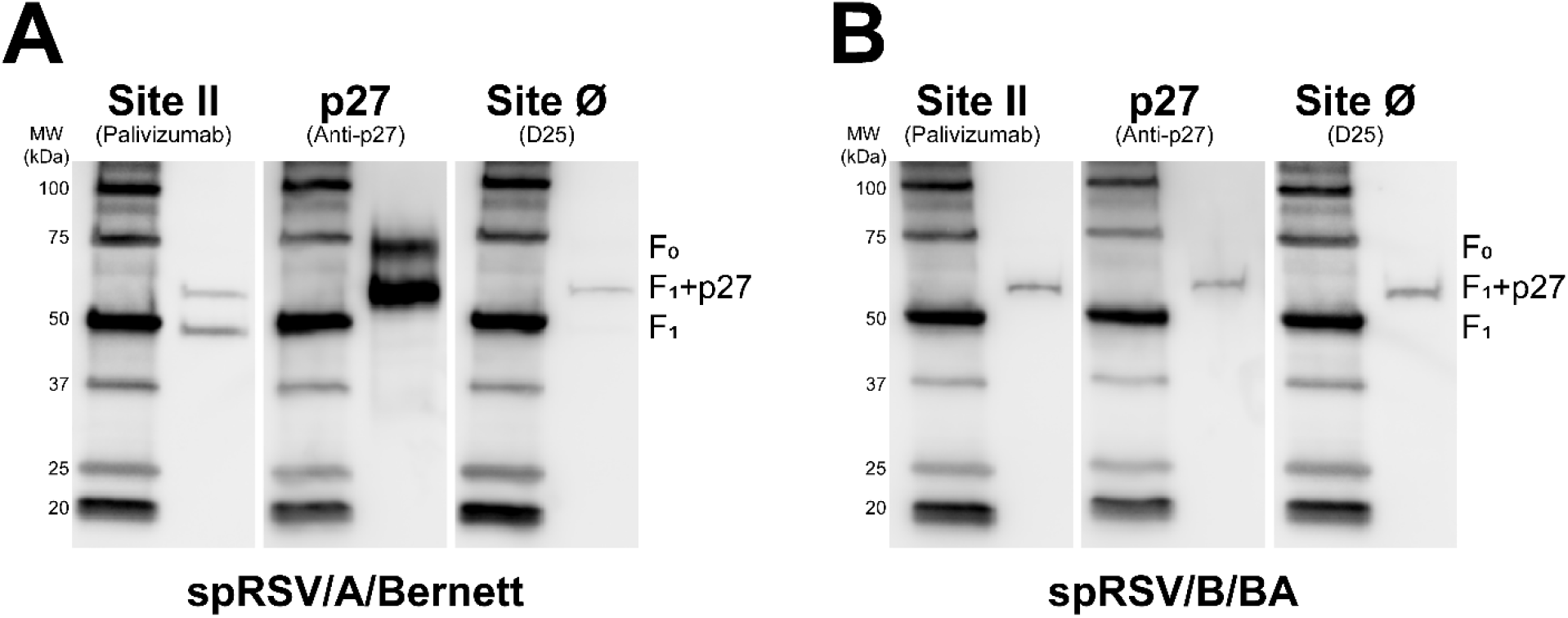
Peptide p27 is present in the F protein of infectious sucrose purified RSV (spRSV) A and B. Equal amounts of spRSV were resolved by reducing SDS-PAGE gel and analyzed by Western blot. (A) spRSV/A/Bernet (B) spRSV/B/BA. Membranes were probed with Palivizumab (Anti-Site II), RSV7.10 (Anti-p27), or D25 (Anti-Site Ø) monoclonal antibodies. F0: uncleaved F protein (∼70 kDa); F1+p27: partially cleaved p27 on the F1 subunit (∼60 kDa); F1: fully cleaved F1 subunit (∼50 kDa).

### Relative quantification of p27 and Site Ø on the F protein of sucrose-purified RSV

We estimated both the relative amount of F protein containing p27 and the F protein in the pre-F conformation by ELISA in spRSV/A/Bernett and spRSV/B/BA. The relative concentration was determined by calculating the ratios of the area under the curve (AUC) of the signal intensity generated by p27 or Site Ø (D25 mAb) over the AUC signal intensity of Site II (Palivizumab mAb) of serially diluted spRSV. Total F protein was estimated with the AUC from the Palivizumab intensity as this mAb binds Site II present in both the pre-F and post-F conformations.

The AUC ratio for p27 detected in spRSV/A/Bernett (Fig. 2A) was 22.5% and for Site Ø it was 34.5% indicating that approximately 22.5% and 34.5% of the F proteins in spRSV/A/Bernett contained p27 and were in a pre-F conformation, respectively. Lower values were detected for the F protein of RSV/B/BA with AUC ratios of 13.7% and 11.9% of F proteins containing p27 and Site Ø, respectively (Fig. 2B). Based on the AUC ratios, spRSV/A/Bernett appeared to have approximately 1.6 times higher proportion of F protein containing p27, and approximately 2.9 times higher proportion of F protein in a pre-F conformation (Site Ø) compared to spRSV/B/BA.

**Figure 2:**
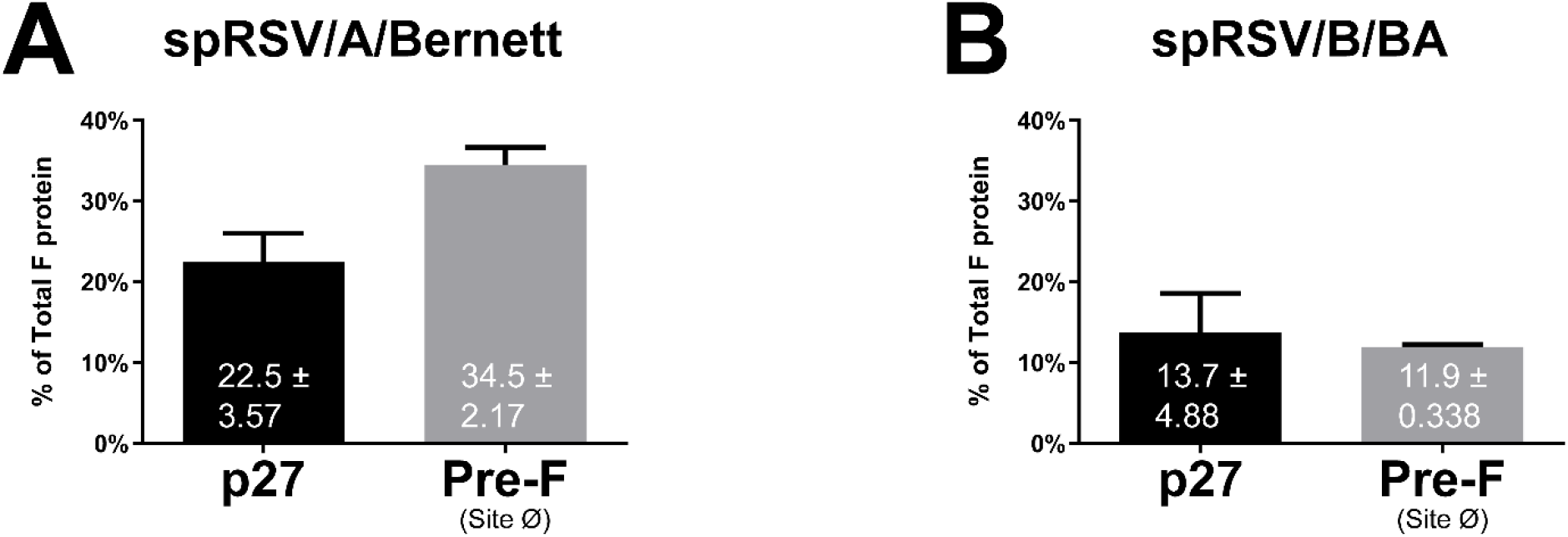
Peptide p27 and F protein in the Pre-Fusion conformation can be detected at quantifiable amounts on the surface of spRSVs. (A) spRSV/A/Bernet (B) spRSV/B/BA (error bars are standard deviation, n=4 replicates). Indirect ELISA of serially diluted spRSVs probed with Palivizumab (Anti-Site II), RSV7.10 (Anti-p27), or D25 (Anti-Site Ø) monoclonal antibodies. Percentages of p27 and Site Ø of the total F protein (Site II) were determined by ratios of Area Under the Curve (AUC) calculated from the OD450 signal from each mAb.

### F protein cleavage and Site Ø stability in RSV-infected HEp-2 and A549 cells measured by flow cytometry

We had previously observed differences in the kinetics of RSV infection by RSV strain and cell type (44), therefore, we speculated that the efficiency of p27 cleavage and the stability of the pre-F conformation may also be RSV strain and cell type dependent. For these studies we used Imaging Flow Cytometry to measure p27, Site Ø and Site II on virus-infected cells; we mitigated the contribution of fluorescence signal from any non-specific intracellular staining using masks to isolate signals to the surface of the cell membrane during data analysis. We used three prototypic strains (RSV/A/Tracy, RSV/A/Bernett and RSV/B/18537) and two contemporary strains (RSV/A/ON and RSV/B/BA) to infect HEp-2 or A549 cell lines.

At each dpi, the same number of events was collected for analysis across all five RSV strains and we determined the percentage of cells that co-stained positive to Site Ø and Site II, or p27 and Site II to monitor the progression of infection and antigenic site expression in HEp-2 and A549 RSV infected cells (Fig. 3).

**Figure 3:**
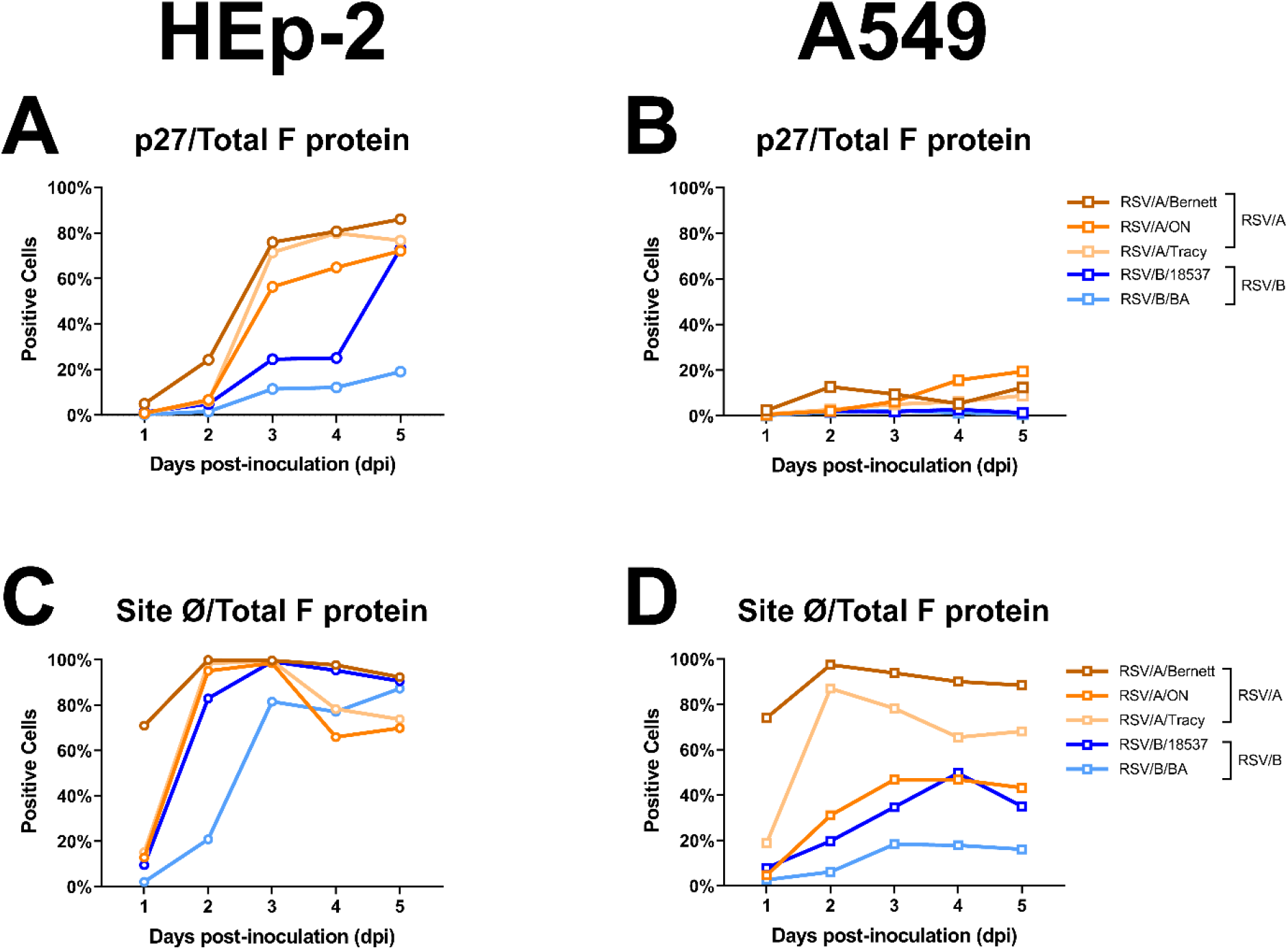
Progression of infection and F protein cleavage are different between HEp-2 and A549 cell lines infected with RSV/A or RSV/B. Imaging Flow Cytometry (Amnis) of HEp-2 cells (A and C) or A549 cells (B and D) infected with RSV/A/Bernett, RSV/A/ON, RSV/A/Tracy, RSV/B/18537, or RSV/B/BA at MOI=0.07. Cells were co-stained with fluorescent-conjugated mAbs (Palivizumab (Anti-Site II), RSV7.10 (Anti-p27), or D25 (Anti-Site Ø)). Percentages of gated double-positive events (p27+Site II+ or Site Ø+Site II+) per day-post-inoculation. Positive populations were determined using uninfected cells.

On 1-dpi, virtually no double-positive cells from RSV infected HEp-2 cell line were positive to p27 (Fig. 3A). At 3-dpi, between 60% and 80% HEp-2 cells infected with three different RSV/A strains exhibited double-positive cells with F proteins containing p27 (Fig. 3A); HEp-2 cells infected with RSV/B/BA reached a maximum of approximately 18% of double-positive cells with detectable p27. Levels of double-positive cells containing p27 on RSV/B/18537 infected HEp-2 cells were low through 4-dpi, but spiked to 63% at 5-dpi, reaching similar levels to the RSV/A strains.

RSV-infected A549 cells also showed differences in double-positive cells with detectable p27 between the RSV/A and the RSV/B strains (Fig. 3B). Unlike the RSV-infected HEp-2 cells, the RSV/A infected A549 cells exhibited a maximum of approximately 20% of double-positive cells presenting F proteins containing p27, and no more than 3% of RSV/B infected A549 cells generated double-positive cells with F proteins containing p27.

Higher levels of detectable p27 on either HEp-2 or A549 cells infected with the three RSV/A strains suggest that the cleavage efficiency of F proteins is lower in RSV/A strains than with the two RSV/B strains. Similarly, lower levels of double-positive cells with detectable p27 in RSV-infected A549 cells suggest that the cleavage efficiency of F proteins is higher in virus-infected A549 cells compared to virus-infected HEp-2 cells.

The number of double-positive cells expressing F proteins with detectable Site Ø (pre-F specific epitope) was different between HEp-2 and A549 cells (Fig. 3C and 3D). On RSV-infected HEp-2 cells (Fig. 3C), at 1-dpi, the percent of double-positive cells presenting F proteins with detectable Site Ø (pre-F conformation) was below 20% for all RSV strains, except for RSV/A/Bernett that had approximately 70% of F proteins with Site Ø. Between 2- and 3-dpi, all three RSV/A strains and RSV/B/18537 reached the maximum percent of double-positive cells expressing F proteins containing Site Ø. The percent of double-positive cells with detectable Site Ø slowly declined to approximately 80% by 5-dpi for the three RSV/A and RSV/B/18537, although it remained relatively stable for RSV/B/BA.

A549 infected cells showed a wider percent distribution of double-positive cells containing F proteins with detectable Site Ø than virus infected HEp-2 cells (Fig. 3D). Of the five RSV strains tested, RSV/A/Bernett and RSV/A/Tracy had the highest percent of double-positive A549 cells with F protein containing Site Ø peaking at 2-dpi. The percent of RSV/A/ON and RSV/B/18537 infected double-positive A549 cells with F protein containing detectable Site Ø peaked at approximately 45%, between 3- and 4-dpi. A549 cells infected with RSV/B/BA had the lowest percent of double-positive cells expressing F proteins containing Site Ø, less than 20% throughout the five days of infection.

### Temperature stress test to evaluate the stability of pre-F conformation in infectious sucrose-purified RSV and RSV-infected cells

The F protein of RSV must maintain its pre-F quaternary structure to fuse with cell membranes. Elevated temperatures can irreversibly trigger the F protein conformation from pre-F (fusogenic) to post-F (non-fusogenic) (5, 6). We evaluated the effect of a temperature-induced conformational change on the F protein by monitoring the stability of Site Ø and changes in p27 detection by ELISA and Imaging Flow Cytometry.

#### F protein conformation on the surface of sucrose-purified RSV virions

For spRSV/A/Bernett and spRSV/B/BA, the ELISA OD signals generated with the mAbs against p27, Site Ø (exclusive to pre-F) and Site II (present in pre-F and post-F) are presented for the varying temperature stress challenges (Supplemental Fig. 2). A transition in the conformation of F protein from pre-F to post-F was observed at 60°C for both spRSV/A/Bernett and spRSV/B/BA, demonstrated by a decrease in mAb binding to Site Ø (Supplemental Fig. 2). Interestingly, an increase in mAb binding was detected against p27 and Site II when evaluating spRSV/A/Bernett. A similar increase in binding was detected for the mAb to Site II in spRSV/B/BA, but not against p27 in the temperature stress experiment (Supplemental Fig. 2). To provide additional insight into the changing OD signals with increasing temperature challenges, the ratio of the OD signal fold-change from 25⁰C for p27 to Site II and Site Ø to Site II were calculated (Fig. 4). A clear distinction was observed for the p27:Site II fold-change ratio between spRSV/A/Bernett (Fig. 4A) and spRSV/B/BA (Fig. 4B). The p27:Site II fold-change ratio remained relatively stable for spRSV/A/Bernett while it dropped to approximately 20% for spRSV/B/BA. Also, with increasing temperature, the Site Ø:Site II fold-change ratio dropped to approximately 20% for spRSV/A/Bernett and near 0% for spRSV/B/BA.

**Figure 4:**
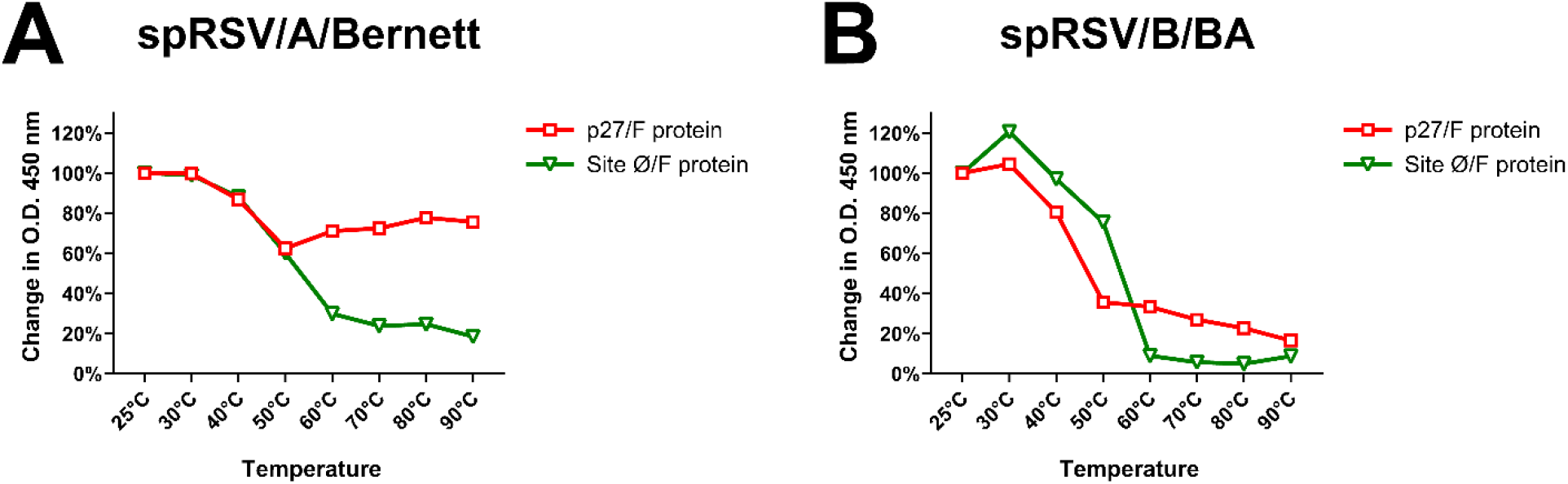
F proteins on spRSV harboring partially cleaved p27 have higher stability of the Pre-Fusion conformation. The percentage change in O.D. 450 nm of the ratios p27/Site II (red squares) or Site Ø/Site II (green triangles) after temperature-induced conformation change of the F protein. ELISA assay of (A) spRSV/A/Bernett, (B) spRSV/B/BA.

#### F protein conformation on the surface of RSV-infected cells

To further investigate the differences we observed between spRSV/A/Bernett and spRSV/B/BA, we used Imaging Flow Cytometry to detect p27, Site II and Site Ø on the F protein expressed on the surface of virus infected cells. We infected HEp-2 cells – the cell line used to generate the sucrose purified RSV strains – as well as A549 cells to determine if host factors also contributed to the stability of the pre-F conformation under different temperature stress challenge. The virus infected cells were harvested at 3-dpi, and increasing temperatures were applied to stress the system prior to staining and fixation. The number of Site II-positive cells was comparable across all temperature points, cell lines, and RSV subtypes (Supplemental Fig. 3), confirming the thermostability of Site II of the F protein in all conditions tested.

On the surface of RSV/A/Bernett infected HEp-2 cells (Fig. 5A and Supplemental Fig. 3A), at 36°C, virtually all cells expressing F protein had mAb binding to Site Ø, consistent with F proteins in a pre-F conformation. Similarly, approximately 90% of the same cells had mAb binding to p27 peptide suggesting most infected cells expressed either uncleaved or partially cleaved F proteins on the cell surface. With increasing temperature, the percentage of RSV/A/Bernett infected HEp-2 cells with mAb binding to p27 and Site Ø declined (Fig. 5A and Supplemental Fig. 3A). At the highest temperature treatment, 65°C, about 40% of HEp-2 cells infected with RSV/A/Bernett had F proteins with detectable Site Ø and p27.

**Figure 5:**
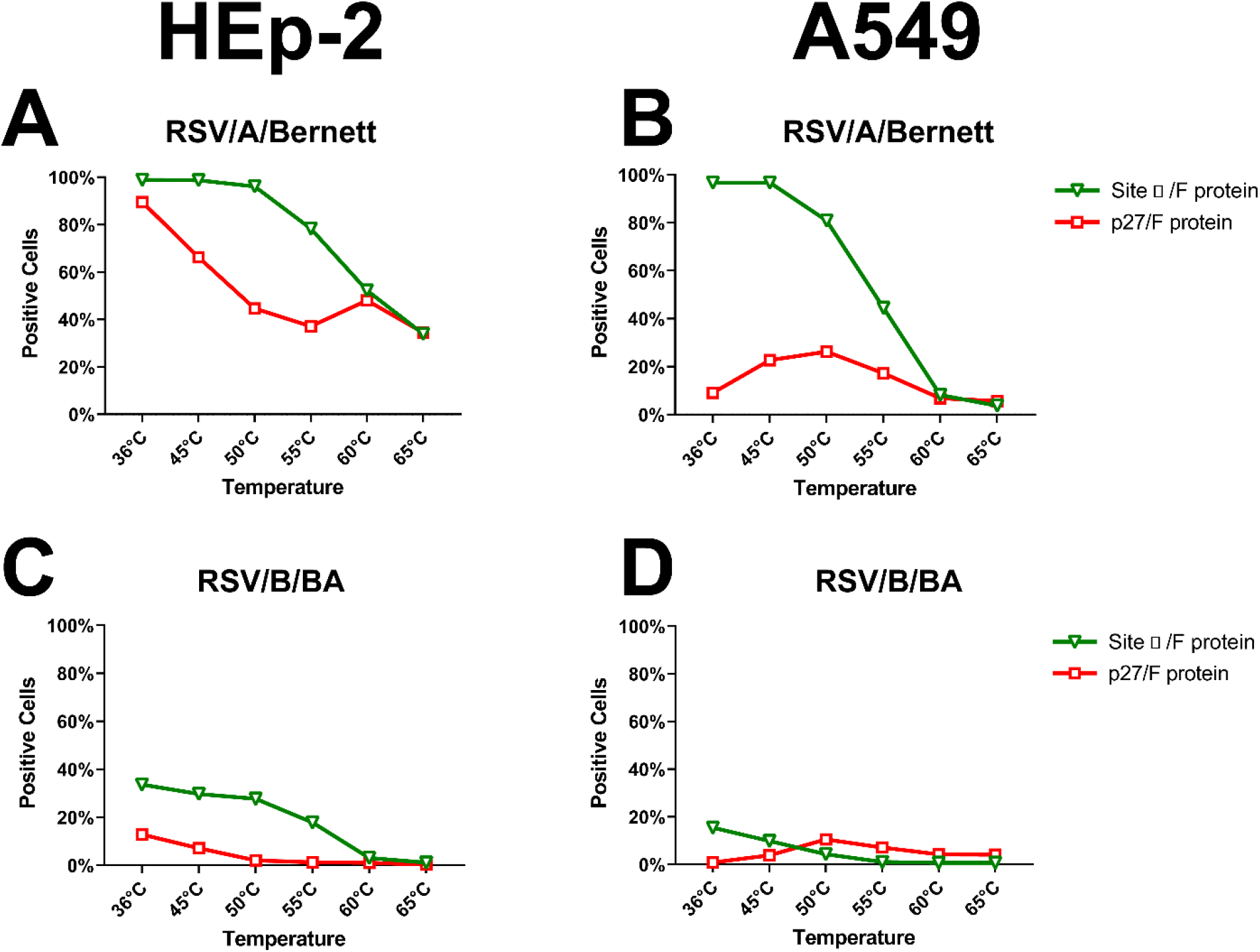
RSV F proteins on the surface of infected cells show presence of partially cleaved p27 has higher stability of the Pre-Fusion conformation. Percentage change in cell count of double-positive populations p27+Site II+ (red squares) and Site Ø+Site II+ (green triangles) after temperature-induced conformational change on RSV-infected cells (MOI=0.07). Imaging Flow Cytometry analysis of (A) HEp-2 cells infected with RSV/A/Bernett; (B) HEp-2 cells infected with RSV/B/BA; (C) A549 cells infected with RSV/A/Bernett; (D) A549 cells infected with RSV/B/BA.

On the surface of A549 cells infected with RSV/A/Bernett (Fig. 5B and Supplemental Fig. 3B), virtually all F proteins had detectable Site Ø at 36°C, but p27 was detected in less than 10% of those cells. With increasing temperatures, mAb binding to Site Ø decreased, indicating transition to the post-F conformation. Interestingly, cells treated at 50°C we observed an increase in mAb binding to p27, reaching a maximum of 23% of the A549 cells infected with RSV/A/Bernett. Further temperature elevation progressively decreased mAb binding to Site Ø and p27, and at 65°C less than 5% of the infected cells had detectable pre-F conformation or p27 peptide.

The F protein on the cell surface of RSV/B/BA infected HEp-2 or A549 cells (Fig. 5C and 5D, and Supplemental Fig. 3C and 3D) behaved in a similar manner to the F protein of RSV/A/Bernett under the same stress conditions, except that the percentage of cells expressing F proteins with detectable p27 and Site Ø was substantially lower. On the surface of RSV/B/BA infected HEp-2 and A549 cells (Fig. 5C and D), at 36°C, 40% and 20% of the cells expressing F proteins, respectively, had detectable Site Ø. P27 was detected in less than 15% of HEp-2 cells infected with RSV/B/BA at 36⁰C (Fig. 5C), and was below 5% on A549 cells (Fig. 5D). Again, an increase in mAb binding to p27 was detected in RSV/B/BA infected A549 cells at 50°C followed by decline to baseline. With increasing temperature stress, the number or percent of cells expressing F proteins with detectable Site Ø and p27 decreased to nearly non-detectable levels in both cell lines infected with RSV/B/BA (Fig. 5C and 5D).

Clear differences were observed in the detection of Site Ø and p27 on the F protein between RSV/A/Bernett and RSV/B/BA, and the stability of these antigenic sites to a temperature stress challenge.

## DISCUSSION

The F protein is well-conserved between RSV/A and RSV/B and is immunogenic, making it the lead candidate for vaccine development. RSV is unique among the negative-sense, single-stranded RNA viruses in that its F protein contains two furin cleavage sites. The F protein requires complete cleavage at both furin cleavage sites, with release of p27, to be fully activated and induce viral fusion to the host cell membrane. The biological role of p27 in the F protein of RSV is largely unknown despite having the greatest amino acid variability between RSV/A and RSV/B strains and containing two or three of the five N-linked glycosylation sites on the F protein (16). Although it is now known that the F protein can also exist partially cleaved, retaining p27 on the surface of RSV/A-infected cells (26), the efficiency of p27 cleavage in different RSV subtypes and genotypes is unknown. To address this knowledge gap, we evaluated the presence of p27 on prototypic and contemporary strains of RSV/A and RSV/B using infectious spRSV and virus-infected HEp-2 and A549 cells. We evaluated the presence of p27 with a mAb that had comparable binding activity against p27 for RSV/A and RSV/B strains (25). We used the Western blot, ELISA, and imaging flow cytometry as orthogonal assays to verify p27 on the F protein. In this report, we demonstrated that p27 was detected in sucrose-purified virions, where spRSV/A/Bernett showed 1.6 and 2.9 times more p27 and Site Ø than spRSV/B/BA. Western blot revealed subgroup differences in the species of F protein, whereby F0 and F1 were both detected in spRSV/A/Bernett while only F1 was detected in spRSV/B/BA, implying different processing of F protein. P27 was preferentially cleaved on the FCS-2 Site of F protein and remained on the N-terminal of the F1 subunit. *In vitro* studies of HEp-2 and A549 cells infected with three prototypic and two contemporary RSV strains demonstrated that the p27 cleavage appeared less efficient on the F protein of RSV/A than RSV/B strains. Lastly, temperature-stress challenges of both spRSV and RSV-infected HEp-2 or A549 cells showed that RSV strains with a higher percent of double-positive cells containing F protein and p27 had greater stability of the metastable pre-F conformation. Altogether, our data suggest that F protein containing p27 has an important role in RSV infection.

RSV F protein has a distinct epitope topology between pre-F and post-F conformations (33, 47). Monoclonal antibodies unique to pre-F conformation (Sites Ø, III, V) and those that target shared sites (sites II and IV) between pre-F and post-F forms are often used to monitor the F conformation (47). For this study, we chose D25 and Palivizumab as the mAbs that target Site Ø and Site II, respectively. F protein binding by both D25 and Palivizumab is consistent with a pre-F form, while F protein binding by only Palivizumab is consistent with a post-F form. We then used a temperature stress test to induce a conformational change from the pre-F to post-F form of the F protein. As expected, binding by the mAb D25 decreased significantly with elevated temperatures. This decrease in D25 binding was not due to denaturation of the F protein because Palivizumab did not lose any binding activity with elevated temperatures. Our premise is supported by a prior thermostability study with uncleaved and cleaved forms of F proteins(45). Ruiz-Arguello et al. used mAbs that bind to site II and site IV and demonstrated the F protein resistance to denaturation when heated up to 100 ºC, after which mAb binding was significantly reduced(45). Thus, the decrease in D25 binding was due to a loss of Site Ø as the F protein transitions to a post-F form.

Elevated temperatures between 45 to 55 ºC can trigger a metastable pre-F conformation to a stable post-F form (46). Interestingly, during the temperature-stress test, we observed an increase in Palivizumab binding activity with spRSV/A/Bernett and RSV/A/BA. An increase in p27 binding activity was also noted but only for spRSV/A/Bernett. We attributed this increase to the opening of the trimeric form during increasing temperature for spRSV/A/Bernett. The “breathing” of the F trimer was recently detected by Gilman et al. Their group showed that the F protein from the A2 strain transitions between monomeric and trimeric folding states, like a “breathing” structure (34). The shift in conformation from pre- to post-F is paramount for the fusogenic activity of the RSV F protein. As we observed differences in levels of p27 and Site Ø between spRSV/A/Bernett and spRSV/B/BA, as well as between HEp-2 and A549-infected cells, we investigated if there was a relationship between p27 and pre-F conformation. Surprisingly, temperature-stress challenges of both spRSV and RSV infected HEp-2, or A549 cells, showed that RSV F proteins with a higher percent of p27 levels had greater stability of the metastable pre-F conformation and thereby likely stabilizing pre-F.

*In vitro* studies, HEp-2 and A549 cells infected with three prototypic and two contemporary RSV strains demonstrated that RSV/As can cause cytopathic effects more rapidly than RSV/Bs, but HEp-2 cells were affected more aggressively than A549 cells (Supplemental Figure 4). These cell lines also differ in their post-translational modification and processing of the F protein, as higher measurable levels of p27 on HEp-2 cells indicated that this cell line was less efficient in cleaving p27 than A549 cells, confirming that HEp-2 and A549 cells elicit different responses to RSV infection. Our previous study demonstrated that HEp-2 cells produce higher cytokine anti-inflammatory responses to RSV infections, while A549 increases the gene expression of antiviral pathways (44). In addition to cell lines, we also demonstrated differences in p27 between RSV strains in RSV/A and RSV/B subgroups. P27 was detected in sucrose-purified virions of RSV/A/Bernett and RSV/B/BA, preferentially cleaved on the FCS-2 Site of F protein, as p27 remained on the N-term of the F1 subunit. Western blot revealed subgroup differences in the species of F protein cleavage products, whereby F0 and F1 were both detected in spRSV/A/Bernett while only F1 was detected in spRSV/B/BA, implying different processing of F protein. It will be important to confirm the presence of p27 in clinical samples collected during RSV infection in children and adults.

To expand the scope of this study, infectivity studies will be necessary to investigate the effects of partially cleaved p27 on infection mechanisms, such as infection with RSV strains lacking the p27 cleavage sites. As we observed differences in the F protein conformation between HEp-2 and A549 infected cells, it would be valuable to later include spRSV strains produced in A549 cells. The publication of a second p27-specific antibody by Lee et al. (26) expands the library of new tools to study p27. Moreover, our results encourage further investigation of the potential role of p27 on RSV infection in more biologically relevant systems other than cancer cell line monolayers, such as nose and lung human organoids, as they better resemble the complexity of the native tissue (48, 49).

Despite the high sequence similarity of the F protein between RSV/As and RSV/Bs, here we reported noticeable differences in behavior and conformation *in vitro*. Additionally, RSV infection in HEp-2 or A549 cell lines produces different innate immune responses and F proteins that differ in their epitope composition and behavior. The F protein from prototypic and contemporaneous RSV strains also showed different quaternary structure dynamics *in vitro*. Previous reports indicated that RSV/A could lead to greater severity of illness in children (50, 51) and generate a higher antibody response in adults (23, 25) than RSV/B. In addition, we observed higher titers of anti-RSV/A p27 serum IgG antibodies than anti-RSV/B p27 IgG antibodies in RSV-infected adults during an RSV/B predominant season (23), possibly explaining the observation that higher levels of p27 are expressed in RSV/A as compared RSV/B. These observations, coupled with the role of the partially cleaved pre-F protein containing p27 in human infection need to be studied with a potential for improved viral fitness. Finally, new therapeutics and vaccines based on the RSV F protein should consider that F proteins from RSV/As or RSV/Bs may have different epitope stability and quaternary conformation bias depending on the cell line it is expressed.

## ACKNOWLEDGMENTS

We want to thank Dr. Gale Smith (Novavax, MD) for kindly providing the RSV7.10 anti-p27 antibody, and the Texas Children’s Hospital William T. Shearer Center for Human Immunobiology for their generous support for this research. This project was supported by the Cytometry and Cell Sorting Core at Baylor College of Medicine with funding from the CPRIT Core Facility Support Award (CPRIT-RP180672), the NIH (CA125123 and RR024574) and the assistance of Joel M. Sederstrom.

## FIGURES AND LEGENDS

**Supplemental Figure 1:**
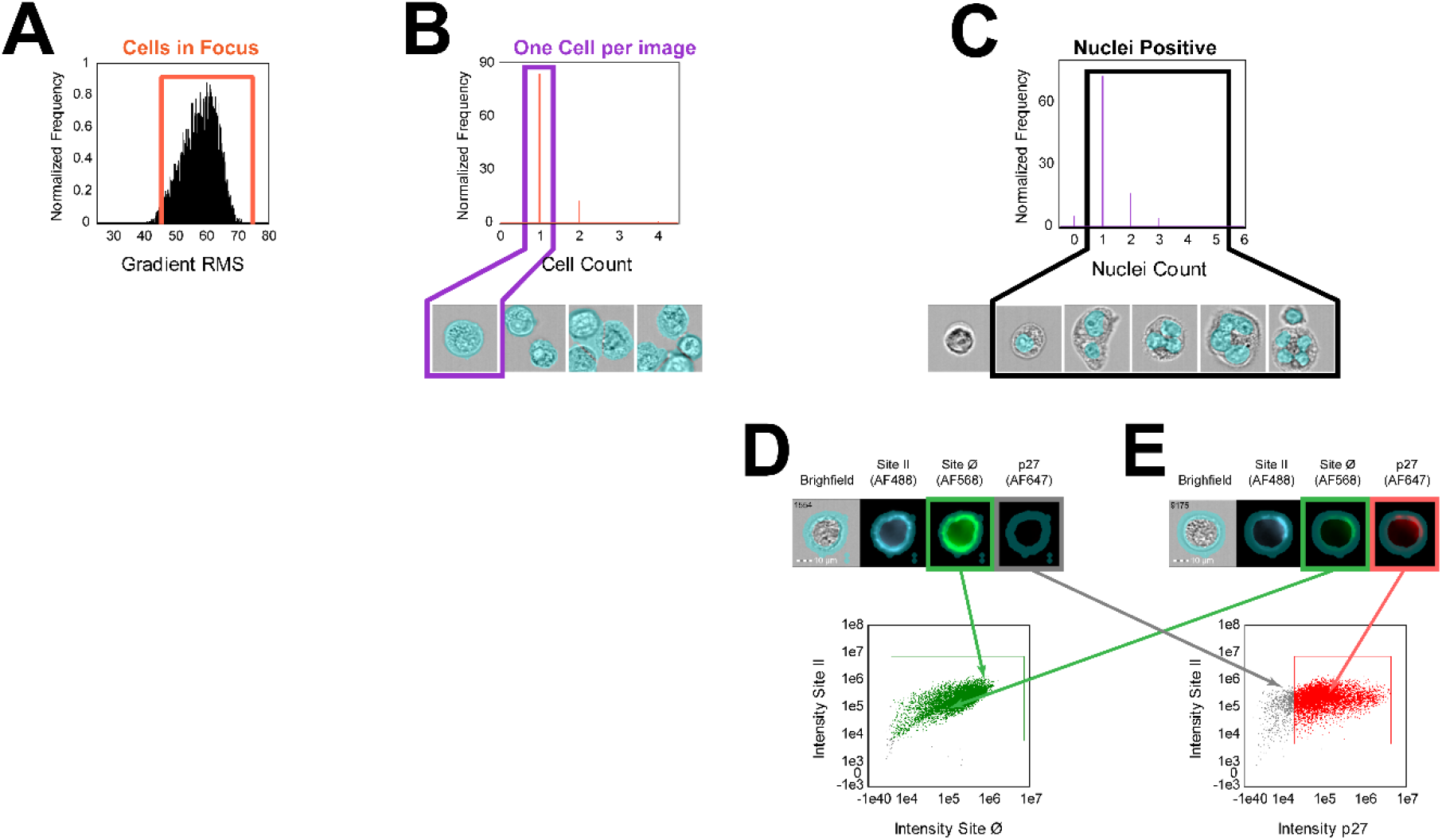
**(A)** Focused cells were gated using gradient RMS feature which helps to select in-focus images with RMS values above 45. **(B)** A Cell Count mask was used to select the images containing only one cell per field of view. **(C)** A Nuclei Count mask determined the number of nuclei per cell and helped highlighting the formation of syncytia following RSV infection *in vitro*. **(D)** and **(E)** Representative images of two RSV-infected cells simultaneously stained for Site II (AF488), Site Ø (AF568), and p27 (AF647). The cyan overlay visible in the brightfield channel is the Cell Membrane mask used to restrict the analysis to the surface of the cell. RSV-infected cells (Site II-positive) with F proteins on the pre-Fusion conformation (Site Ø-positive, green arrow), **(D)** without p27 (p27-negative, grey arrow) or **(E)** with p27 (p27-positive, red arrow). **(F)** Dot plot of Site Ø-positive cells in Site II-positive RSV-infected cells (green dots). Dot plot of gated p27-positive cells in Site II-positive RSV-infected cells (red dots). All cell images were captured with 40 × objective at default flow speed in PBS. Scale bar = 10 µm. Mask formulas: • Cell Count mask: Range(Watershed(Object(M01, Ch01, Tight)), 250-5000, 0-1), • Nuclei Count mask: Intensity(Morphology(M07, Ch07), Ch07, 800-4095), • Cell Membrane mask: Dilate(M01, 3) And Not Erode(M01, 8).

**Supplemental Figure 2:**
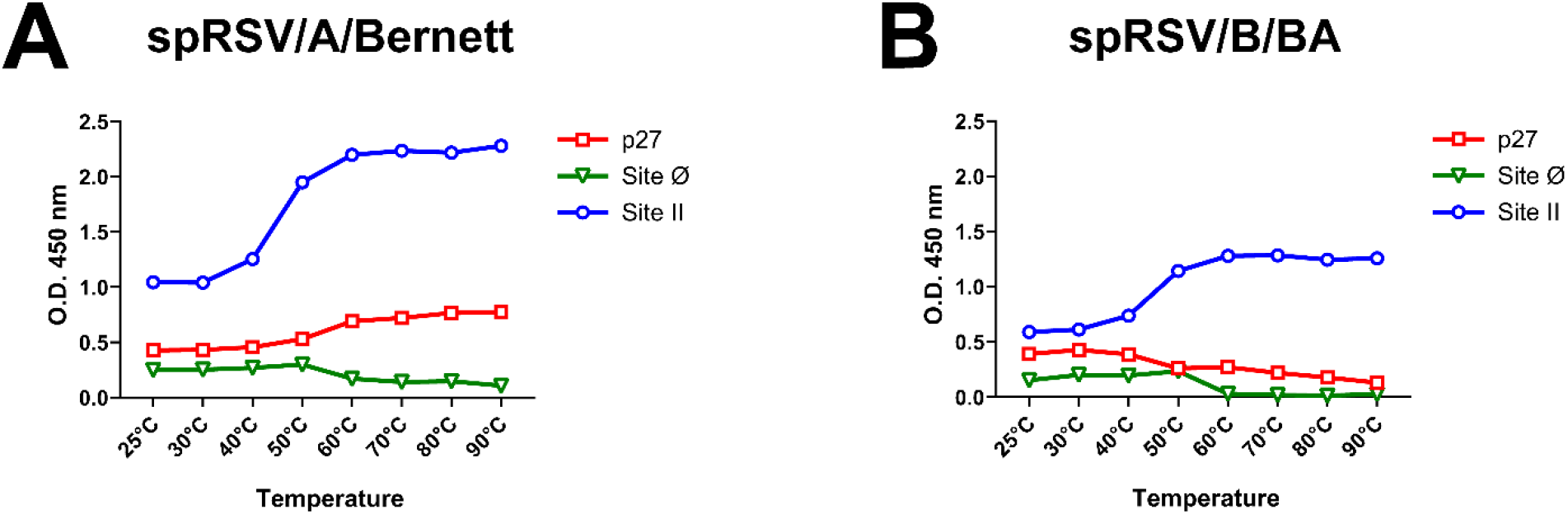
Changes in O.D. 450 nm of antigenic Site II (blue circles), Site Ø (green triangles) and p27 (red squares) (A) spRSV/A/Bernett or (B) spRSV/B/BA by ELISA.

**Supplemental Figure 3:**
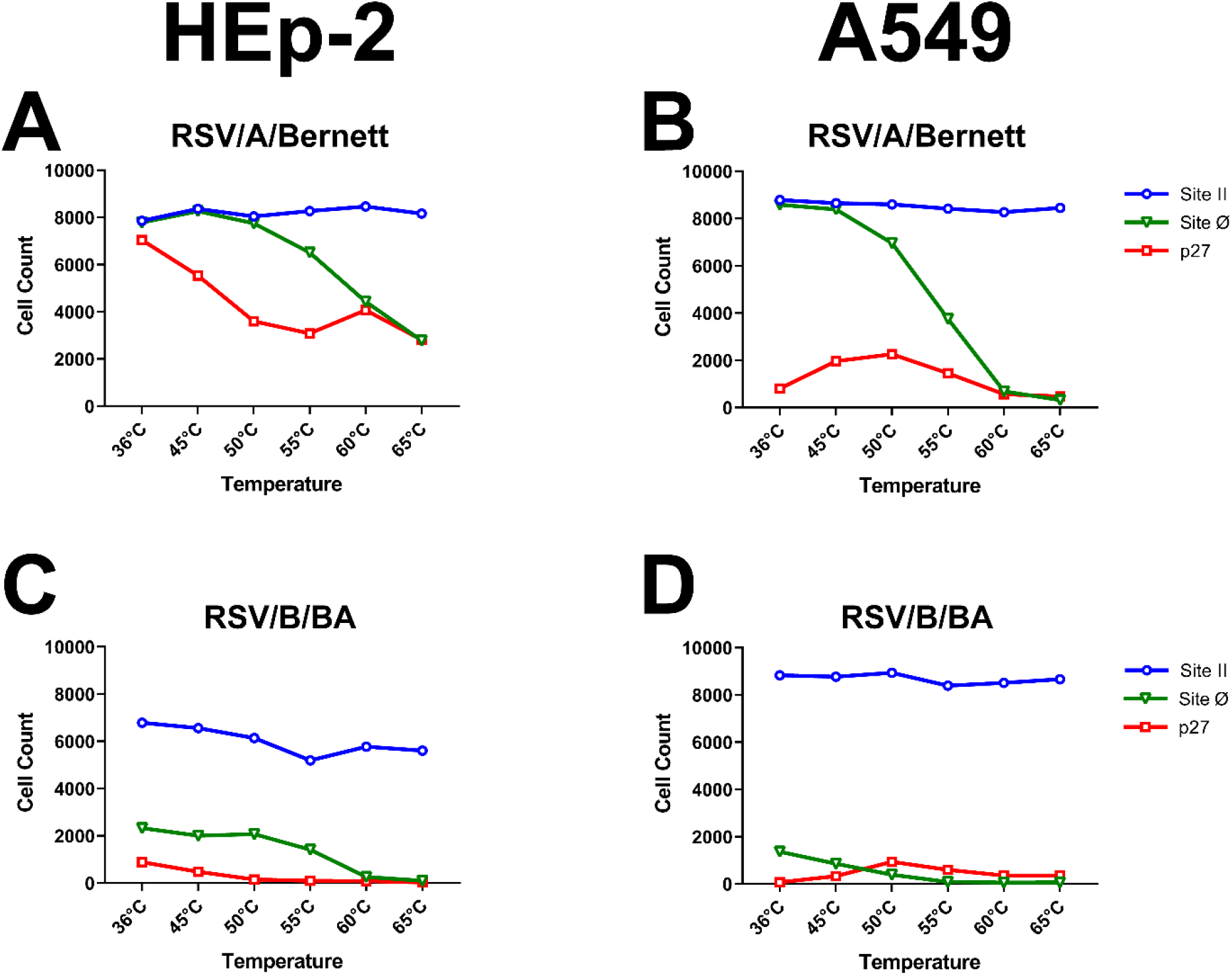
Measuring changes in antigenic site detection and stability during a temperature-stress experiment on RSV-infected HEp-2 (A and C) and A549 (B and D) cells. Cell Count of cells positive for Sites II (blue circles), Site Ø (green triangles) and p27 (red squares). Cells infected with MOI=0.07 of RSV/A/Bernett and RSV/B/BA (Buenos Aires). Cells were treated at 36, 45, 50, 55, 60 or 65°C for 10 minutes, cooled down to room temperature and stained for Imaging Flow Cytometry data collection.

**Supplemental Figure 4:**
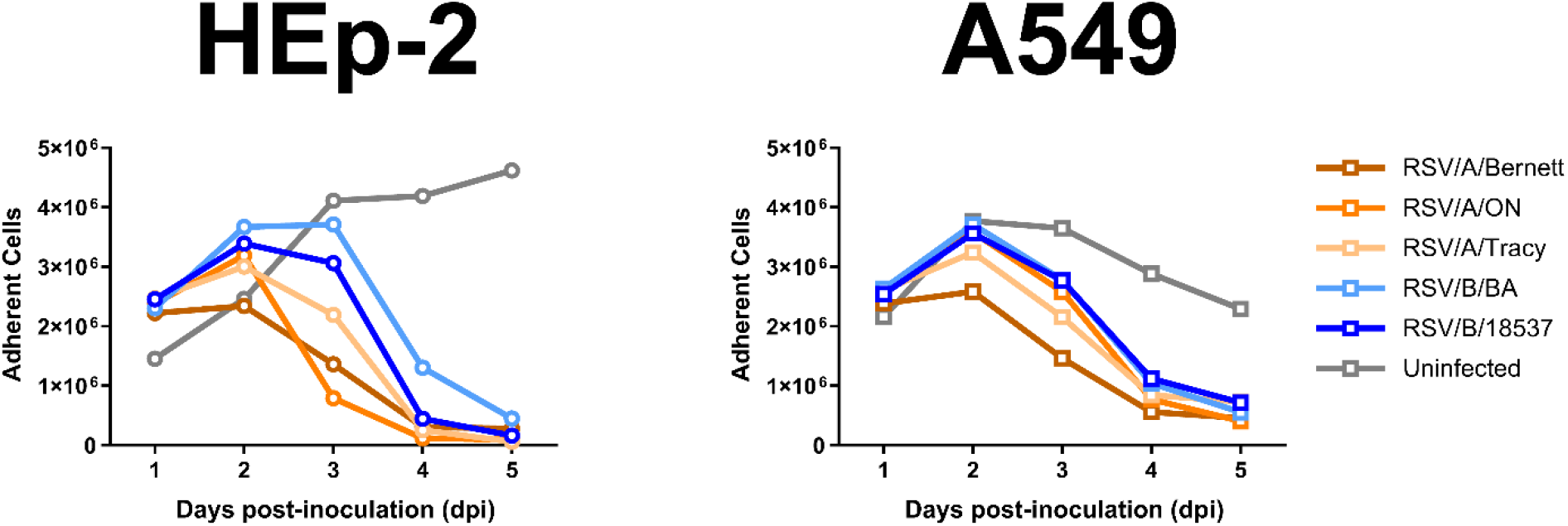
Progression of cell detachment of HEp-2 and A549 cells infected with various RSVs at days 1 to 5 post-inoculation, starting at day 1 (MOI=0.07). Uninfected cells (grey circles) were used as control for cell attachment. At each timepoint, culture media was removed, the cells were carefully rinsed with PBS, and the remaining attached cells lifted with 1X Versene solution. After a PBS wash, cells were centrifuged at 350 xg for 6 minutes, the volume was brought up to 1 mL of PBS and counted using a Cellometer Mini Automated Cell Counter (Nexcelom Biosciences).

## REFERENCES

1. Shi T, McAllister DA, O’Brien KL, Simoes EAF, Madhi SA, Gessner BD, Polack FP, Balsells E, Acacio S, Aguayo C, Alassani I, Ali A, Antonio M, Awasthi S, Awori JO, Azziz-Baumgartner E, Baggett HC, Baillie VL, Balmaseda A, Barahona A, Basnet S, Bassat Q, Basualdo W, Bigogo G, Bont L, Breiman RF, Brooks WA, Broor S, Bruce N, Bruden D, Buchy P, Campbell S, Carosone-Link P, Chadha M, Chipeta J, Chou M, Clara W, Cohen C, de Cuellar E, Dang D-A, Dash-yandag B, Deloria-Knoll M, Dherani M, Eap T, Ebruke BE, Echavarria M, de Freitas Lázaro Emediato CC, Fasce RA, Feikin DR, Feng L, Gentile A, Gordon A, Goswami D, Goyet S, Groome M, Halasa N, Hirve S, Homaira N, Howie SRC, Jara J, Jroundi I, Kartasasmita CB, Khuri-Bulos N, Kotloff KL, Krishnan A, Libster R, Lopez O, Lucero MG, Lucion F, Lupisan SP, Marcone DN, McCracken JP, Mejia M, Moisi JC, Montgomery JM, Moore DP, Moraleda C, Moyes J, Munywoki P, Mutyara K, Nicol MP, Nokes DJ, Nymadawa P, da Costa Oliveira MT, Oshitani H, Pandey N, Paranhos-Baccalà G, Phillips LN, Picot VS, Rahman M, Rakoto-Andrianarivelo M, Rasmussen ZA, Rath BA, Robinson A, Romero C, Russomando G, Salimi V, Sawatwong P, Scheltema N, Schweiger B, Scott JAG, Seidenberg P, Shen K, Singleton R, Sotomayor V, Strand TA, Sutanto A, Sylla M, Tapia MD, Thamthitiwat S, Thomas ED, Tokarz R, Turner C, Venter M, Waicharoen S, Wang J, Watthanaworawit W, Yoshida L-M, Yu H, Zar HJ, Campbell H, Nair H. 2017. Global, regional, and national disease burden estimates of acute lower respiratory infections due to respiratory syncytial virus in young children in 2015: a systematic review and modelling study. The Lancet 390:946–958.

2. Mazur NI, Löwensteyn YN, Willemsen JE, Gill CJ, Forman L, Mwananyanda LM, Blau DM, Breiman RF, Madhi SA, Mahtab S, Gurley ES, el Arifeen S, Assefa N, Scott JAG, Onyango D, Tippet Barr BA, Kotloff KL, Sow SO, Mandomando I, Ogbuanu I, Jambai A, Bassat Q, Thamthitiwat S, Gentile A, Lucion MF, Pires MR, De-Paris F, Gordon A, Sánchez JF, Lucero MG, Lupisan SP, Gessner BD, Tall H, Halasa N, Khuri-Bulos N, Nokes DJ, Munywoki PK, Otieno GP, O’Brien KL, Oshitani KL, da Costa Oliveira MT, de Freitas Lázaro Emediato CC, Ali A, Aamir UB, Noyola DE, Cohen C, Moyes J, Giamberardino HIG, Webler JM, de Matos Bezerra PG, do Bezerra Duarte MCM, Chu HY, Das RR, Weber MW, Homaira N, Jaffe A, Sturm-Ramirez KM, Su W, Yuan CC, Chaves S, Emukule GO, de Andrade Nishioka S, de Carvalho FC, Gökçe Ş, Raboni SM, Hawkes M, Messaoudi M, Bryant J, Dbaibo GS, Hanna-Wakim R, Sampath Jayaweera Jaa, Stolyarov K, Suntarattiwong P, Mussá T, Bruno A, de Mora D, Wanlapakorn N, de Xie Z, Ai J, Ojeda J, Zamora L, Obodai E, Odoom JK, Ismail MT, Buchwald A, O’Callaghan-Gordo C, Fernandez-Sarmiento J, Obando-Belalcazar E, Dhole T, Verma S, Eski A, Kartal GO, al Amad M, al Serouri AW, Funchan Y, Sam JIC, Jarovsky D, da Silva Dgbp, Perales JG, Toh TH, Yit JLS, Kendirli T, Gun E, Sagna T, Diagbouga S, Chowdhury F, Islam MA, Venter M, Visser A, Pham MH, Vásquez-Hoyos P, González-Dambrauskas S, Rubio FD, Karsies T, Zemanate E, Izquierdo L, Palomino RL, Pardo-Carrero R, Grigolli-Cesar R, Menta S, Monteverde N, Duyu M, Saha S, Saha SK, Kelly M, Echavarria M, Tran T, Borgi A, Ayari A, Caballero MT, Polack FP, Omer S, Kazi AM, Simões EAF, Satav A, Bont LJ. 2021. Global Respiratory Syncytial Virus-Related Infant Community Deaths. Clinical Infectious Diseases 73:S229–S237.

3. Li Y, Reeves RM, Wang X, Bassat Q, Brooks WA, Cohen C, Moore DP, Nunes M, Rath B, Campbell H, Nair H, RSV Global Epidemiology Network S, RESCEU investigators WJ, Antonio M, Talavera GA, Badarch D, Baillie VL, Barrera-Badillo G, Bigogo G, Broor S, Bruden D, Buchy P, Byass P, Chipeta J, Clara W, Dang D-A, Emediato CC de FL, Jong M de, Díaz-Quiñonez JA, Do LAH, Fasce RA, Feng L, Ferson MJ, Gentile A, Gessner BD, Goswami D, Goyet S, Grijalva CG, Halasa N, Hellferscee O, Hessong D, Homaira N, Jara J, Kahn K, Khuri-Bulos N, Kotloff KL, Lanata CF, Lopez O, Bolaños MRL, Lucero MG, Lucion F, Lupisan SP, Madhi SA, Mekgoe O, Moraleda C, Moyes J, Mulholland K, Munywoki PK, Naby F, Nguyen TH, Nicol MP, Nokes DJ, Noyola DE, Onozuka D, Palani N, Poovorawan Y, Rahman M, Ramaekers K, Romero C, Schlaudecker EP, Schweiger B, Seidenberg P, Simoes EAF, Singleton R, Sistla S, Sturm-Ramirez K, Suntronwong N, Sutanto A, Tapia MD, Thamthitiwat S, Thongpan I, Tillekeratne G, Tinoco YO, Treurnicht FK, Turner C, Turner P, Doorn R van, Ranst M van, Visseaux B, Waicharoen S, Wang J, Yoshida L-M, Zar HJ. 2019. Global patterns in monthly activity of influenza virus, respiratory syncytial virus, parainfluenza virus, and metapneumovirus: a systematic analysis. Lancet Glob Health 7:e1031–e1045.

4. Walsh EE. 2017. Respiratory Syncytial Virus Infection: An Illness for All Ages. Clin Chest Med 38:29–36.

5. Glezen WP, Taber LH, Frank AL, Kasel JA. 1986. Risk of primary infection and reinfection with respiratory syncytial virus. Am J Dis Child 140:543–6.

6. Ohuma EO, Okiro EA, Ochola R, Sande CJ, Cane PA, Medley GF, Bottomley C, Nokes DJ. 2012. The Natural History of Respiratory Syncytial Virus in a Birth Cohort: The Influence of Age and Previous Infection on Reinfection and Disease. Am J Epidemiol 176:794–802.

7. Curran D Eliazar |, Cabrera S, Bracke B, Raymond K, Foster A, Umanzor C Goulet | Philibert, Powers Iii JH. 2022. Impact of respiratory syncytial virus disease on quality of life in adults aged ≥50 years: A qualitative patient experience cross-sectional study. Influenza Other Respir Viruses https://doi.org/10.1111/IRV.12929.

8. Falsey AR, Walsh EE. 2005. Respiratory Syncytial Virus Infection in Elderly Adults. Drugs Aging 22:577–587.

9. Hall CB, Walsh EE, Long CE, Schnabel KC. 1991. Immunity to and Frequency of Reinfection with Respiratory Syncytial Virus. J Infect Dis 163:693–698.

10. Tang JW, Loh TP. 2014. Correlations between climate factors and incidence—a contributor to RSV seasonality. Rev Med Virol 24:15–34.

11. Peret TC, Hall CB, Hammond GW, Piedra PA, Storch GA, Sullender WM, Tsou C, Anderson LJ. 2000. Circulation patterns of group A and B human respiratory syncytial virus genotypes in 5 communities in North America. J Infect Dis 181:1891–6.

12. Tabor DE, Fernandes F, Langedijk AC, Wilkins D, Lebbink RJ, Tovchigrechko A, Ruzin A, Kragten-Tabatabaie L, Jin H, Esser MT, Bont LJ, Abram ME. 2021. Global molecular epidemiology of respiratory syncytial virus from the 2017-2018 INFORM-RSV study. J Clin Microbiol 59.

13. Obando-Pacheco P, Justicia-Grande AJ, Rivero-Calle I, Rodríguez-Tenreiro C, Sly P, Ramilo O, Mejías A, Baraldi E, Papadopoulos NG, Nair H, Nunes MC, Kragten-Tabatabaie L, Heikkinen T, Greenough A, Stein RT, Manzoni P, Bont L, Martinón-Torres F. 2018. Respiratory Syncytial Virus Seasonality: A Global Overview. J Infect Dis 217:1356–1364.

14. Pandya MC, Callahan SM, Savchenko KG, Stobart CC. 2019. A contemporary view of respiratory syncytial virus (RSV) biology and strain-specific differences. Pathogens 8:1–15.

15. Battles MB, McLellan JS. 2019. Respiratory syncytial virus entry and how to block it. Nat Rev Microbiol 17:233–245.

16. Hause AM, Henke DM, Avadhanula V, Shaw CA, Tapia LI, Piedra PA. 2017. Sequence variability of the respiratory syncytial virus (RSV) fusion gene among contemporary and historical genotypes of RSV/A and RSV/B. PLoS One 12:1–21.

17. Tan L, Coenjaerts FEJ, Houspie L, Viveen MC, van Bleek GM, Wiertz Ejhj, Martin DP, Lemey P. 2013. The Comparative Genomics of Human Respiratory Syncytial Virus Subgroups A and B: Genetic Variability and Molecular Evolutionary Dynamics. J Virol 87:8213–8226.

18. Zimmer G, Budz L, Herrler G. 2001. Proteolytic Activation of Respiratory Syncytial Virus Fusion Protein. Journal of Biological Chemistry 276:31642–31650.

19. Gonzalez-Reyes L, Ruiz-Arguello MB, Garcia-Barreno B, Calder L, Lopez JA, Albar JP, Skehel JJ, Wiley DC, Melero JA. 2001. Cleavage of the human respiratory syncytial virus fusion protein at two distinct sites is required for activation of membrane fusion. Proceedings of the National Academy of Sciences 98:9859–9864.

20. Chang A, Dutch RE. 2012. Paramyxovirus fusion and entry: Multiple Paths to a common end. Viruses 4:613–636.

21. Zimmer G, Conzelmann K-K, Herrler G. 2002. Cleavage at the Furin Consensus Sequence RAR/KR 109 and Presence of the Intervening Peptide of the Respiratory Syncytial Virus Fusion Protein Are Dispensable for Virus Replication in Cell Culture. J Virol 76:9218–9224.

22. Krzyzaniak MA, Zumstein MT, Gerez JA, Picotti P, Helenius A. 2013. Host Cell Entry of Respiratory Syncytial Virus Involves Macropinocytosis Followed by Proteolytic Activation of the F Protein. PLoS Pathog 9.

23. Blunck BN, Aideyan L, Ye X, Avadhanula V, Ferlic-Stark L, Zechiedrich L, Gilbert BE, Piedra PA. 2022. Antibody responses of healthy adults to the p27 peptide of respiratory syncytial virus fusion protein. Vaccine 40:536–543.

24. Fuentes S, Coyle EM, Beeler J, Golding H, Khurana S. 2016. Antigenic Fingerprinting following Primary RSV Infection in Young Children Identifies Novel Antigenic Sites and Reveals Unlinked Evolution of Human Antibody Repertoires to Fusion and Attachment Glycoproteins. PLoS Pathog 12:1–24.

25. Ye X, de Rezende WC, Iwuchukwu OP, Avadhanula V, Ferlic-Stark LL, Patel KD, Piedra FA, Shah DP, Chemaly RF, Piedra PA. 2020. Antibody response to the furin cleavable twenty-seven amino acid peptide (P27) of the fusion protein in respiratory syncytial virus (RSV) infected adult hematopoietic cell transplant (HCT) recipients. Vaccines (Basel) 8:192.

26. Lee J, Lee Y, Klenow L, Coyle EM, Tang J, Ravichandran S, Golding H, Khurana S. 2022. Protective antigenic sites identified in respiratory syncytial virus fusion protein reveals importance of p27 domain. EMBO Mol Med 14:1–14.

27. Swanson KA, Settembre EC, Shaw CA, Dey AK, Rappuoli R, Mandl CW, Dormitzer PR, Carfi A. 2011. Structural basis for immunization with postfusion respiratory syncytial virus fusion F glycoprotein (RSV F) to elicit high neutralizing antibody titers. Proceedings of the National Academy of Sciences 108:9619–9624.

28. McLellan JS, Yang Y, Graham BS, Kwong PD. 2011. Structure of Respiratory Syncytial Virus Fusion Glycoprotein in the Postfusion Conformation Reveals Preservation of Neutralizing Epitopes. J Virol 85:7788–7796.

29. McLellan JS, Chen M, Joyce MG, Sastry M, Stewart-Jones GBE, Yang Y, Zhang B, Chen L, Srivatsan S, Zheng A, Zhou T, Graepel KW, Kumar A, Moin S, Boyington JC, Chuang G-Y, Soto C, Baxa U, Bakker AQ, Spits H, Beaumont T, Zheng Z, Xia N, Ko S-Y, Todd J-P, Rao S, Graham BS, Kwong PD. 2013. Structure-based design of a fusion glycoprotein vaccine for respiratory syncytial virus. Science 342:592–8.

30. Xie Q, Wang Z, Ni F, Chen X, Ma J, Patel N, Lu H, Liu Y, Tian JH, Flyer D, Massare MJ, Ellingsworth L, Glenn G, Smith G, Wang Q. 2019. Structure basis of neutralization by a novel site II/IV antibody against respiratory syncytial virus fusion protein. PLoS One 14:1–20.

31. Krarup A, Truan D, Furmanova-Hollenstein P, Bogaert L, Bouchier P, Bisschop IJM, Widjojoatmodjo MN, Zahn R, Schuitemaker H, McLellan JS, Langedijk JPM. 2015. A highly stable prefusion RSV F vaccine derived from structural analysis of the fusion mechanism. Nat Commun 6:8143.

32. Swanson KA, Balabanis K, Xie Y, Aggarwal Y, Palomo C, Mas V, Metrick C, Yang H, Shaw CA, Melero JA, Dormitzer PR, Carfi A. 2014. A Monomeric Uncleaved Respiratory Syncytial Virus F Antigen Retains Prefusion-Specific Neutralizing Epitopes https://doi.org/10.1128/JVI.01225-14.

33. McLellan JS, Chen M, Leung S, Graepel KW, D. X, Yang Y, Zhou T, Baxa U, Yasuda E, Beaumont T, Kumar A, Modjarrad K, Zheng Z, Zhao M, Xia N, Kwong PD, Graham BS. 2013. Structure of RSV Fusion Glycoprotein Trimer Bound to a Prefusion-Specific Neutralizing Antibody. Science (1979) 340:1113–1117.

34. Gilman MSA, Furmanova-Hollenstein P, Pascual G, B. van w‘t Wout A, Langedijk JPM, McLellan JS. 2019. Transient opening of trimeric prefusion RSV F proteins. Nat Commun 10:2105.

35. McLellan JS, Ray WC, Peeples ME. 2013. Structure and Function of Respiratory Syncytial Virus Surface Glycoproteins, p. 83–104. In Current Topics in Microbiology and Immunology.

36. Liljeroos L, Krzyzaniak MA, Helenius A, Butcher SJ. 2013. Architecture of respiratory syncytial virus revealed by electron cryotomography. Proc Natl Acad Sci U S A 110:11133–11138.

37. Farzan SF, Palermo LM, Yokoyama CC, Orefice G, Fornabaio M, Sarkar A, Kellogg GE, Greengard O, Porotto M, Moscona A. 2011. Premature activation of the paramyxovirus fusion protein before target cell attachment with corruption of the viral fusion machinery. Journal of Biological Chemistry 286:37945–37954.

38. Patel N, Massare MJ, Tian JH, Guebre-Xabier M, Lu H, Zhou H, Maynard E, Scott D, Ellingsworth L, Glenn G, Smith G. 2019. Respiratory syncytial virus prefusogenic fusion (F) protein nanoparticle vaccine: Structure, antigenic profile, immunogenicity, and protection. Vaccine 37:6112–6124.

39. Melero JA, Mas V, McLellan JS. 2017. Structural, antigenic and immunogenic features of respiratory syncytial virus glycoproteins relevant for vaccine development. Vaccine 35:461–468.

40. Piedra PA, Cron SG, Jewell A, Hamblett N, McBride R, Palacio MA, Ginsberg R, Oermann CM, Hiatt PW, McColley S, Bowman M, Borowitz D, Castile R, McCoy K, Prestige C, Brown ME, Stevens J, Regelmann W, Milla C, Sammut P, Colombo J, Eisenberg J, Murphy TD, Finder J, Kurland G, Winnie G, Orenstein D, Voter K, Light M, Pian MS, Harris C, Stokes D, Fink R, Ren C, Gorvoy J, Varlotta L, Dyson M. 2003. Immunogenicity of a new purified fusion protein vaccine to respiratory syncytial virus: a multi-center trial in children with cystic fibrosis. Vaccine 21:2448–2460.

41. Piedra PA, Glezen WP, Kasel JA, Welliver RC, Jewel AM, Rayford Y, Hogerman DA, Hildreth SW, Paradiso PR. 1995. Safety and immunogenicity of the PFP vaccine against respiratory syncytial virus (RSV): the Western blot assay aids in distinguishing immune responses of the PFP vaccine from RSV infection. Vaccine 13:1095–1101.

42. Kim HW, Canchola JG, Brandt CD, Pyles G, Chanock RM, Jensen K, Parrott RH. 1969. Respiratory syncytial virus disease in infants despite prior administration of antigenic inactivated vaccine. Am J Epidemiol 89:422–34.

43. Ye X, Iwuchukwu OP, Avadhanula V, Aideyan LO, McBride TJ, Ferlic-Stark LL, Patel KD, Piedra F-A, Shah DP, Chemaly RF, Piedra PA. 2018. Comparison of Palivizumab-Like Antibody Binding to Different Conformations of the RSV F Protein in RSV-Infected Adult Hematopoietic Cell Transplant Recipients. J Infect Dis 217:1247–1256.

44. Rajan A, Piedra F-A, Aideyan L, McBride T, Robertson M, Johnson HL, Aloisio GM, Henke D, Coarfa C, Stossi F, Menon VK, Doddapaneni H, Muzny DM, Cregeen SJJ, Hoffman KL, Petrosino J, Gibbs RA, Avadhanula V, Piedra PA. 2021. Multiple RSV strains infecting HEp-2 and A549 cells reveal cell line-dependent differences in resistance to RSV infection. bioRxiv 2021.06.15.448622.

45. Ruiz-Argüello MB, Martín D, Wharton SA, Calder LJ, Martín SR, Cano O, Calero M, García-Barreno B, Skehel JJ, Melero JA. 2004. Thermostability of the human respiratory syncytial virus fusion protein before and after activation: implications for the membrane-fusion mechanism. Journal of General Virology 85:3677–3687.

46. Yunus AS, Jackson TP, Crisafi K, Burimski I, Kilgore NR, Zoumplis D, Allaway GP, Wild CT, Salzwedel K. 2010. Elevated temperature triggers human respiratory syncytial virus F protein six-helix bundle formation. Virology 396:226–237.

47. McLellan JS. 2015. Neutralizing epitopes on the respiratory syncytial virus fusion glycoprotein. Curr Opin Virol 11:70–75.

48. Rijsbergen LC, Lamers MM, Comvalius AD, Koutstaal RW, Schipper D, Duprex WP, Haagmans BL, de Vries RD, de Swart RL. 2021. Human Respiratory Syncytial Virus Subgroup A and B Infections in Nasal, Bronchial, Small-Airway, and Organoid-Derived Respiratory Cultures. mSphere 6.

49. Porotto M, Ferren M, Chen YW, Siu Y, Makhsous N, Rima B, Briese T, Greninger AL, Snoeck HW, Moscona A. 2019. Authentic modeling of human respiratory virus infection in human pluripotent stem cell-derived lung organoids. mBio 10.

50. Laham FR, Mansbach JM, Piedra PA, Hasegawa K, Sullivan AF, Espinola JA, Camargo CA. 2017. Clinical Profiles of Respiratory Syncytial Virus Subtypes A and B among Children Hospitalized with Bronchiolitis. Pediatr Infect Dis J 36:808.

51. Papadopoulos NG, Gourgiotis D, Javadyan A, Bossios A, Kallergi K, Psarras S, Tsolia MN, Kafetzis D. 2004. Does respiratory syncytial virus subtype influences the severity of acute bronchiolitis in hospitalized infants? Respir Med 98:879–882.

